# Biosensors Characterization: Formal methods from the Perspective of Proteome Fractions

**DOI:** 10.1101/2023.12.01.569588

**Authors:** Nicolás A. Vaccari, Dahlin Zevallos-Aliaga, Tom Peeters, Daniel G. Guerra

## Abstract

Many studies characterize transcription factors and other regulatory elements to control the expression of heterologous genes in recombinant systems. However, most lack a formal approach to analyse the parameters and context-specific variations of these regulatory components. This study addresses this gap by establishing formal and convenient methods for characterising regulatory circuits. We model the bacterial cell as a collection of a small number of proteome fractions. Then, we derive the proteome fraction over time and obtain a general theorem describing its change as a function of its expression fraction, which represents a specific portion of the total biosynthesis flux of the cell. Formal deduction reveals that when the proteome fraction reaches a maximum, it becomes equivalent to its expression fraction. This equation enables the reliable measurement of the expression fraction through direct protein quantification. In addition, experimental data demonstrate a linear correlation between protein production rate and specific growth rate over a significant time period. This suggests a constant expression fraction within this window. The expression fractions estimated from the slopes of these intervals and those obtained from maximum protein amount points can both be independently fitted to a Hill function. In the case of an IPTG biosensor, in five cellular contexts, expression fractions determined by the maximum method and the slope method produced similar dose-response parameters. Additionally, by analysing two more biosensors, for mercury and cumate detection, we demonstrate that the slope method can be effectively applied to various systems, generating reliable Hill function parameters.

## Introduction

Synthetic biology relies on the assumption that biological systems can be engineered by design, assembling parts and circuits like modules to create new systems of increasing complexity [1, 2]. However, regulatory circuits and even individual components often exhibit variable behaviours depending on the cellular context or culture conditions [3, 4, 5, 6]. In addition, research groups employ different techniques and data analysis methods when characterising these regulatory elements experimentally. This variation further hinders the ability to exchange parts in a modular approach.

Despite these challenges, the field has grown steadily. New biological parts are constantly being incorporated for novel engineered functions, leading to practical successes at different levels. For example, transcription factors and DNA or RNA sequences can link intracellular metabolites or extracellular signals to gene expression, controlling native or heterologous genes [7, 8, 9]. This strategy is applied in creating new biosynthetic pathways [10, 11], identifying optimal strains for the production of a molecule of interest[12, 13], and developing biosensors for various applications such as environmental monitoring, food safety, and disease diagnosis [14, 15, 16, 17]. Furthermore, the activity of specific pathways can be regulated through transcription factors or sensing RNA elements, thus introducing a dynamic control to optimise metabolic flux toward the desired biosynthesis [18, 19]. There are a large number of characterised transcription factors, evaluated by their dose-response, detection limit, sensitivity, and dynamic range [20, 21, 19, 22]. However, the lack of standardised procedures to analyse their ability to regulate reporter genes hinders progress in establishing robust libraries of modular tools. For example, there is no consensus on how to select culture time points that allow reliable measurement averaging between replicates or facilitate comparisons of output under different conditions, cellular contexts, and laboratories [23, 24].

This study addresses this challenge by proposing a theoretical framework for the characterisation of regulatory circuits. This framework is based on the perspective that the cell is composed of a few proteome fractions, a concept that was used to study bacterial metabolism and growth regulation [25, 26, 27, 28, 29, 30, 31, 32]. By mathematically defining the proteome fraction and deriving it over time, we introduce a theorem that links the evolution of a set of proteins to a specific fraction of the cell’s biosynthesis rate, which we call the expression fraction. We formally demonstrate that this expression fraction can be quantified by two alternative methods: directly by analysing a single point of peak accumulation of reporter protein; or through the estimation of a linear relationship between specific production and specific growth rates during a given interval. We tested the reciprocal consistency of these two methods experimentally by comparing the dose-response profile of different *Escherichia coli* strains induced by varying IPTG concentrations and two other biosensors based on transcription factors, for the detection of mercury and cumate.

We consider that the framework presented in this article helps to tackle well-known challenges that are still prevalent in synthetic biology today [3, 33]. Firstly, it offers readily reproducible methodologies for characterising genetic circuits and their components, relying on easily measurable data such as culture growth and reporter protein levels. Secondly, our approach simplifies the mathematical modelling of genetic regulation systems by masking complexity through the concept of proteome fractions, while still capturing the interplay between native and heterologous cellular functions. This facilitates a deeper understanding of the inherent properties of sensing circuits, differentiating them from characteristics that depend on the cell and culture context. Finally, we speculate on how to extend this analysis to investigate orthogonality and potential crosstalk between heterologous and native genetic components.

## Methods

### *E. coli* strains

MG1655 ΔendA ΔrecA (DE3) was a gift from Kristala Prather (Addgene code # 37854) [34]. BLR(DE3) and BL21(DE3) agar stabs were generously provided by Carlos Bustamante at UC Berkeley and Mauricio Baez at the Universidad de Chile, respectively. Cells were rendered competent by the CaCl_2_ incubation protocol and transformed by 42 °C heat shock with pUC-T7GFP or simultaneously with pUC-T7GFP and pACYCDuet-1. Luria broth (LB) selection agar plates were supplemented with glucose 1% to reduce the basal expression of T7 and ampicillin to select cells transformed with the plasmid pUC-T7GFP, or ampicillin and chloramphenicol to select those with the plasmids pUC-T7GFP and pACYCDuet-1.

### Plasmids

Sequences were designed using Benchling RRID:SCR 013955 [Biology Software] (2020). Retrieved from https://benchling.com. Twist Bioscience synthesised a gene block containing the following: T7 promoter, lac operator, Shine-Dalgarno ribosome binding sequence, a codon-optimised GFPmut3 coding sequence [35] fused with the LVA tag for rapid degradation (AANDENYLVA) [36], and the T7 terminator. The complete sequence is available in GenBank, accession number PQ015608. A high copy number GFP expression plasmid, pUC-T7GFP, was built by inserting this construct into the pUC57 vector through standard digestion and ligation reactions between the restriction sites NcoI and KpnI. To introduce additional expression of LacI, cells were transformed with the low-copy plasmid pACYCDuet-1, acquired from GenScript.

The mercury biosensor was constructed by inserting the *merR* gene, which encodes the MerR mercury sensitive transcription factor, and the promoter from transposon Tn501 from *Pseudomonas aeruginosa* plasmid pVS1 (GenBank: Z00027.1), containing the MerR DNA binding site, in the pUC57 vector. This enabled conditional expression of the *RFP* reporter gene in response to mercury. NEB^®^ Stable *E. coli* competent cells were transformed with the resulting plasmid. The complete design, cloning procedures, and characterisation are described in [37].

A cumate biosensor was constructed based on a regulatory circuit adapted from Choi et al. [38]. Briefly, a constitutively expressed CymR protein represses a promoter that controls the expression of GFPmut3 fused with the LVA tag for rapid degradation (AANDENYLVA). Cumate-induced dissociation of CymR activates GFP expression. The circuit sequence was synthesised as a gBlock by Integrated DNA Technologies, Inc. and cloned at the NcoI and NotI sites of the pUC57 vector. The resulting plasmid was transformed into BL21(DE3), BLR(DE3) and MG1655(DE3) competent cells. The complete sequence is available in GenBank, accession number PQ010742.

### Culture conditions

The Specific growth rates and fluorescence data presented here were obtained from 200 *µ*L microcultures in M9 medium supplemented with 0.2% casamino acids and 0.24% glucose, prepared as follows: Freshly transformed colonies were handpicked to inoculate 10 mL of M9 medium supplemented with ampicillin, or ampicillin and chloramphenicol when pACYCDuet-1 was included. These cultures were incubated at 37 ^°^C and 250 RPM overnight, their cell density was measured and harvested by centrifugation at 5000 RPM for 5 minutes. Immediately, cells were resuspended in fresh medium to the appropriate volume to obtain an optical density of 0.1 (approximately 2 × 10^7^ cells·mL^−1^) and were loaded in a 96 microtiter plate. At this point, reporter protein expression was induced in 195 *µ* L of culture by manually pipetting 5 *µ*L of the appropriate inducer (IPTG, HgBr_2_, cumate), reaching the final concentration indicated for each case, completing 200 *µ*L in each well. These microcultures were incubated at 30 ^°^C with oscillatory shaking at 220 RPM for 16 hours without the addition of antibiotics. The microtiter plate was covered with its plastic lid and the outer wells were filled with 200 *µ*L of water so that a similar humidity surrounded all monitored cultures. This configuration resulted in a volume loss of less than 2. 5% due to evaporation after 16 hours.

### Measurements

Culture density (OD_600_) and fluorescence were monitored in a plate reader (Tecan Infinite 200 Pro). The production of GFP was registered with the following settings: 475 nm excitation and 516 nm emission, with the signal gain set at 50 out of 100. Similarly, the production of RFP was recorded using 585-nm excitation and 608-nm emission, with the signal gain set at 95. In both cases, fluorescence data were collected with 25 flashes, reading every 15 minutes.

### Total protein quantification and visualisation

For this assay, two flasks (for each strain), containing 50 mL of transparent medium, were inoculated with the bacteria so that the final OD_600_ was 0.1. One of the flasks was induced with IPTG (0.5 mM) at the beginning of the culture. The cultures were then left to grow at 30 °C. Cultures were sampled at three time points: 0 h, 2 h and 4 h. For each of the samples taken during bacterial growth, two tests were carried out. Quantification of total proteins using the Bradford reagent and visualisation of total proteins using a polyacrylamide gel. For the total protein quantification essay, 2 mL of culture was taken and the bacteria were pelleted at 13500 rpm for 5 min. This pellet was then resuspended in 500 *µ*L of 8M Urea and lysed by three cycles of freeze thawing for 15 min at -70°C and 15 min at 37 ° C. Finally, 10 *µ*L of this suspension was mixed with 300 *µ*L of Bradford reagent (Thermo Fisher Scientific), incubated 10 min at room temperature and measured at 595 nm for protein quantification. For visualisation of the total proteins essay, 2 mL of culture was pelleted, resuspended in 500 *µ*L of 2% SDS and heated at 95 ° C for 10 min. Subsequently, this suspension was run on a 15% polyacrylamide gel.

### Estimation of the heterologous fraction, specific growth rate and specific production rate rate

We considered fluorescence directly proportional to the mass of the reporter protein (GFP or RFP) and the optical density proportional to the total protein mass. Thus, the ratio of fluorescence to optical density, 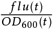, was measured to estimate the heterologous fraction, *φ*_*H*_.

To calculate the specific growth rate at each time point *µ*(*t*), the optical density was measured at *t* and at the immediate next time point *t* + Δ*t*. The difference between both measurements was divided by Δ*t* and the optical density at the former point *t*:

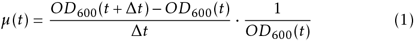

The specific production rate of *GFP* (or *RFP*) was estimated by normalizing the slope of fluorescence accumulation over time. For GFPmut3, the fluorophore maturation time is 4.1 minutes[35], therefore, no maturation delay was introduced in the calculations as it is shorter than our measurement time window of 15 minutes. On the other hand, the merR system used *RFP* as a reporter which has a maturation time of 1 hour, meaning that the red fluorescence that was observed in any given time is proportional to the amount of *RFP* produced 4 measuring points earlier. We corrected this by shifting the fluorescence curve back in time by 4 points. Finally, we calculated *ρ*_*H*_ by measuring the fluorescence at time *t* and at next time point *t* + Δ*t*. We divide the difference by time step Δ*t* and optical density at time *t*:

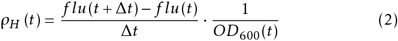

### Calculations

Data handling, processing, plotting, regression analysis, and modeling were performed using the R programming language. All linear regressions were performed using R’s built-in lm and nls functions. The data set employed was uploaded in figshare (DOI: https://doi.org/10.6084/m9.figshare.26314723.v1) [39]. To determine the expression fractions using the maximum method, the time series data for *φ*_*H*_ were grouped by strain and induction condition (N=8 for BL21(DE3) with the T7 system, N=9 for the MerR and CymR systems, and N = 4 for all other cases). Subsequently, 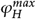 was identified for each series within each group and these values were averaged. The average 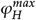 was obtained at various inducer concentrations and then used in the Hill fitting process.

For the slope method, the expression fraction was determined as the slope obtained from a linear regression between the specific growth rate and the specific production rate during growth (*µ* ¿ 0). To establish the appropriate time window for this regression, all specific growth rate data for each strain under each induction condition were analysed. At each time point, a simple t test was used to assess whether the average value of the specific growth rate (N=8 for BL21(DE3) with the T7 system, N=9 for the MerR and CymR systems and N = 4 for all other cases) differed significantly from zero. If a significant difference was found (*α* = 0.05), the time point was considered part of the growth phase and included in the regression analysis. Subsequently, the earliest data points were iteratively excluded one by one until the determination coefficient (*R*^2^) reached a maximum value, which was greater than 0.9 in all cases. The slopes of the regressions obtained at various inducer concentrations were registered and used to fit the Hill function.

Dose-response profiles were used to build Hill functions for each biosensor. A scatterplot was created depicting the expression fractions data (obtained from slopes or 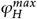 values) versus the inducer concentration for each titration experiment. The nls function with the port algorithm was then used to fit a Hill function to this plot. Lower bounds of 0 were set for the values of *H* (maximum expression) and *k*_*I*_ (affinity constant), while a lower bound of 1 was set for the value of *n* (Hill coefficient). The weight of each data point was determined by the inverse of its variance, which assigned a higher weight to more precise measurements.

## Results

### DE3 system illustrates the challenges of biosensors characterisation

We began our investigation experimenting with the DE3 expression system in B and K-12 *E. coli* strains. This system combines a native element, the LacI repressor, and a heterologous one, the T7 RNA polymerase (T7RNAP)[40], and is widely applied in the laboratory-scale production of recombinant proteins [41]. In addition, the LacI repressor served as a model for biosensor studies [42], and both the LacI repressor and T7RNAP have been used as modular components in the construction of logic gates within synthetic regulatory circuits [43, 44, 45].

For our DE3 system experiments, we used a plasmid containing the GFP gene under the control of the T7 promoter and a LacI operator. Using this setting as an IPTG biosensor, we employed a two-stage culture protocol. The first stage was focused on the accumulation of biomass, then in a subsequent phase, starting at OD_600_ = 0.1, the expression was induced by IPTG addition. This configuration revealed variations in specific growth rates and GFP output both between different *E. coli* strains and between those with and without additional expression of the LacI repressor (introduced through the plasmid pACYCDuet), as shown in Figure 1.

**Figure 1.**
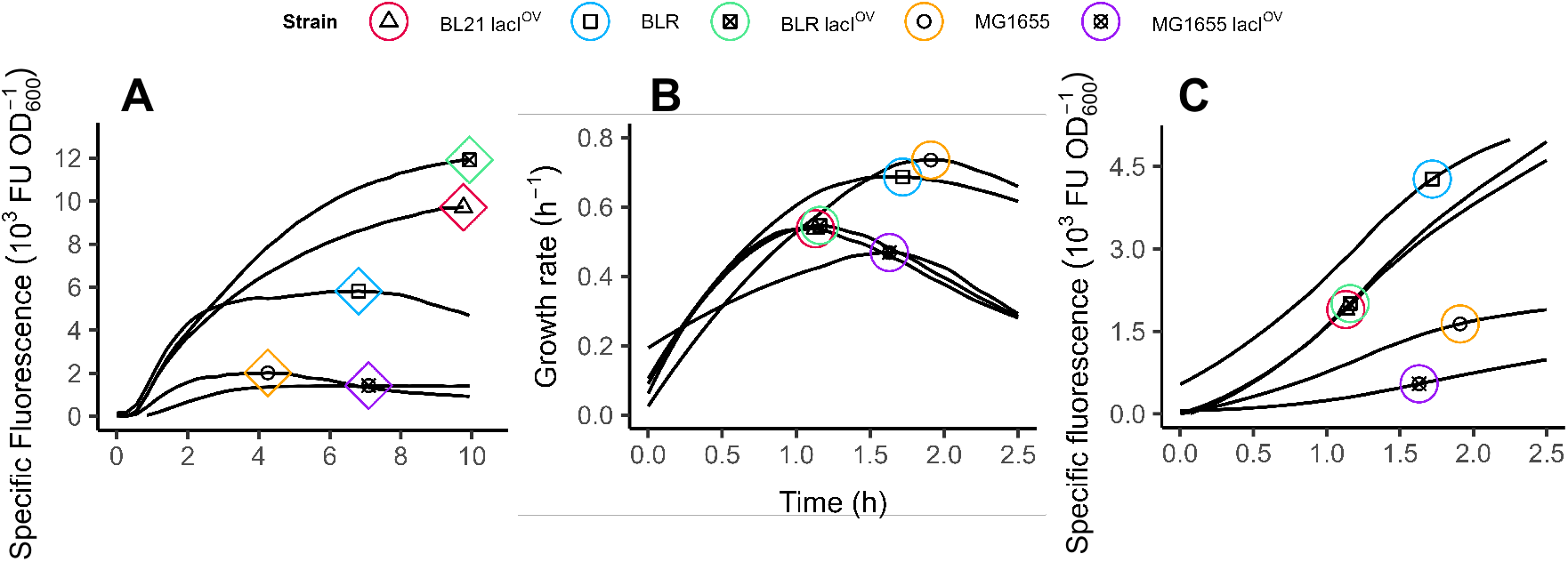
Analysing the heterologous proteome fraction *φ*_*H*_ at different time points. A) The specific fluorescence in time represents the accumulation of GFP over 10 hours of cultivation after induction by 1 mM IPTG. Diamonds indicate the point of maximum specific fluorescence, 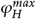. B) Specific growth rate of the same systems; the circles indicate the point of maximum specific growth rate in each culture. C) The specific fluorescence in the same experiment, within the first 2.5 hours. ALT TEXT: images comparing the normalised fluorescence when measured at the moment of peak growth rate vs measuring its maximum value.

Although differences between cultures are visible, analysing them requires applying objective criteria that are not self-evident. A common option to compare biosensors is to analyse their results at the peak of their specific growth rate, which does not necessarily align at any particular time of culture in the case of different strains. In our case, even though all cultures started at the same cellular density and were stimulated with IPTG at the same time, BLR(DE3) and MG1655(DE3) showed their maximum specific growth rates later than BL21(DE3) as shown in Figure 1. Consequently, if these points are selected, the production of GFP by BL21(DE3) and BLR(DE3) with additional LacI would be plotted just after 30 minutes of induction, when the fluorescence is still relatively low, while those of BLR(DE3) and MG1655(DE3) would be evaluated after longer accumulation (Figure 1 C). Although the peak of the specific growth rate is a common reference for metabolism and regulation studies, in our case analysing the protein yield at these time points would lead to unfair comparisons. This misrepresents the different strains and could imply wrong interpretations of the circuit’s properties. For example, strain MG1655(DE3) would appear to be a producer similar to B strains simply because the latter are analysed after a shorter protein accumulation.

This case illustrates that analysing a culture’s response to heterologous expression requires identifying a time point that offers biologically meaningful information, taking into account the induction strategy and the culture’s evolution over time. When aiming for maximum yield or signal, experimenters often choose the stationary phase to analyse the accumulated protein. Alternatively, through trial and error, analysis may be shifted to an earlier time point, as is common in the case of recombinant protein production, where short induction times minimise the potentially undesirable consequences of heterologous overexpression [46, 47, 48], such as toxicity and protein misfolding. In the case of developing biosensors for analyte detection or for dynamically controlling engineered metabolic pathways, it is preferable to centre the analysis during exponential growth, near the peak of the specific growth rate, as this is when the anabolic capacity is at its maximum. In addition, studies of bacterial physiology use the growth peak as a reference because this is when bacterial metabolism reaches an optimum within the constraints imposed by external conditions and internal regulations. However, as stated above, focussing on the time point of maximum specific growth rate might not be informative in the case of comparing different strains in scenarios such as two-stage cultures, consisting of a biomass accumulation phase and a consecutive production phase. Given this circumstance, a new criterion is needed to select measurement points to reliably study genetic circuits, such as biosensors, in a manner that enables accurate comparisons and reproducibility.

### A theorem for the evolution of proteome fractions in time: the concept of expression fraction

We argue that when analysing the production of heterologous proteins in general, and whole cell biosensors in particular, the focus should shift from protein yield to biosynthesis flux allocation. We conceptualised the general biosynthesis rate divided into a set of rates (Figure 2), each producing a different type of protein, termed a proteome fraction (*φ*_*i*_). Similarly as has been done in models previously reported [26, 28, 29, 25, 30, 27, 31], proteins will be grouped together as a unified proteome fraction if they share a similar biological function and are regulated by the same cues [25, 32]. In previous work, a proteome fraction that fulfils the core structural functions will remain constant (*φ*_*Q*_), while the other two fractions are variable and perform nutrient assimilation (*φ*_*C*_ (*t*)) and biomass production (*φ*_*R*_(*t*)), respectively. Finally, in the case of one set of heterologous proteins controlled by a single circuit, these will constitute a fraction called the heterologous fraction (*φ*_*H*_ (*t*)). The total anabolic flux will be divided into the synthesis of specific protein sets, in proportions denoted as the expression fractions, represented by *f*_*i*_ (*t*), where *i* is the protein group to which it is dedicated (*f*_*R*_, *f*_*Q*_, *f*_*H*_, *f*_*C*_ in figure 2D). In this way, the expression fraction *f*_*i*_ can be understood as the proportion of total ribosomal activity that synthesises the proteome fraction *φ*_*i*_. The value of *f*_*i*_ (*t*) will depend on the specific regulatory mechanism of the protein set and the conditions of the cell. The full conceptual and mathematical reasoning behind the expression fraction is detailed in the supplementary material.

**Figure 2.**
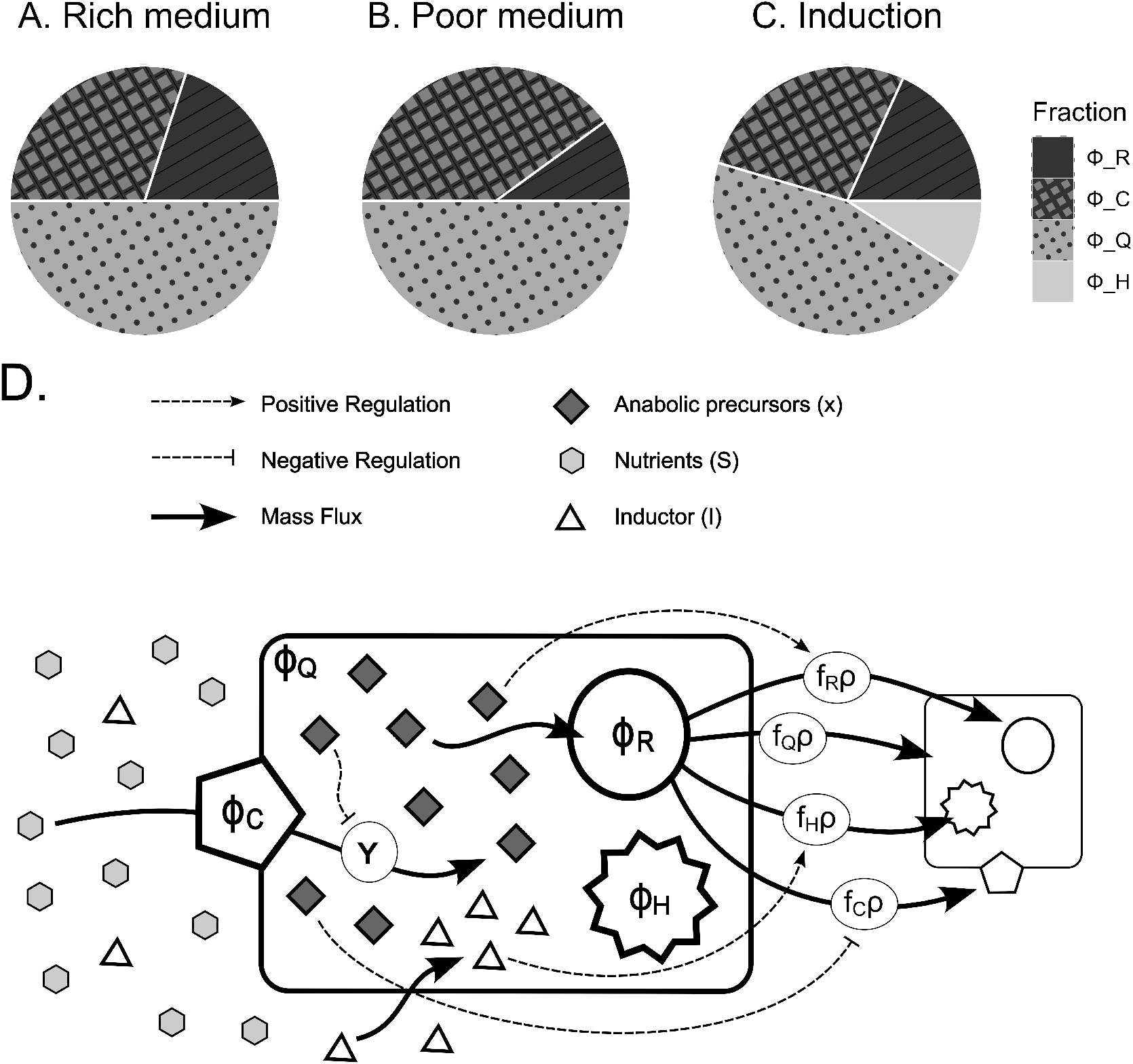
A model of proteome allocation and metabolic rate. Representation of the proteome fractioning of a cell: A) in a rich medium that allows the cell to maximize its specific growth rate through the activity of ribosomal fraction, *φ*_*R*_; B) in a poor medium, where the cell maximizes its fixation of nutrients through the activity of the carbon fixator fraction, *φ*_*C*_ ; C) producing a heterologous fraction, *φ*_*H*_, causing a redistribution of the native proteome allocation. D) Representation of rates within bacteria. Biochemical transformations are represented by solid arrows. The light gray hexagons represent the nutrients *S*(*t*), which are converted into anabolic precursors, *x*(*t*), (dark gray diamonds) by the nutrient fixator, *φ*_*C*_ (pentagon) at a rate *Y*. The anabolic precursors are then used by the ribosomal fraction, *φ*_*R*_ (circle), at a rate *ρ*, to produce new biomass (represented by the small cell on the right side). Biosynthesis is divided into a multi-outlet branching of specific protein production rates (black arrows), each one with their respective expression fractions, *f*_*R*_(*t*), *f*_*C*_ (*t*), *f*_*Q*_ (*t*), *f*_*H*_ (*t*). Dashed arrows and dashed blunt arrows represent positive and negative regulation, respectively. The concentration of anabolic precursors stimulates the specific rate of *φ*_*R*_ production and represses the specific rate of *φ*_*C*_ production; the specific rate of *φ*_*Q*_ production is constant and the specific rate of *φ*_*H*_ production is activated by its specific inducer (triangles), which diffuses from the extracellular space. ALT TEXT: Graphical representation of the proteome fractions and idealisation of the cell’s activity as compound by a set of fluxes for carbon fixation and protein biosynthesis

With these definitions, solving the derivation of a proteome fraction over time (complete derivation in the Biosynthesis Allocation Theorem section of the supplementary material) produces an equation that links the change in a proteome fraction to its expression fraction and the total biosynthesis rate, *ρ*(*t*), as follows:

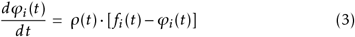

In the subsequent sections, we will explain how *f*_*i*_ (*t*) can be determined from easily obtainable experimental data.

### Finding the expression fraction and its regulation: The maximum method

A remarkable consequence emerging from equation 3 is that for any proteome fraction, when its differential over time equals zero 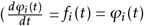, the expression fraction and the proteome fraction must be equal *f*_*i*_ (*t*) = *φ*_*i*_ (*t*) (assuming that the cell’s anabolic rate is never zero as long as it is alive). Therefore, regardless of the growth behaviour of a culture, registering the heterologous fraction when it has reached a maximum should correctly represent the expression of the circuit. Data collected at this point under various conditions can be used reliably to build a function that represents the regulated expression of the circuit, for example, in response to the variation of the inducer concentration.

It is generally accepted that gene expression responds to the concentration of an inducer following a Hill function [49], a model commonly applied to biosensors [42]. Using a Hill equation for representing the regulation of the heterologous fraction as a function of the inducer *I*, we obtain the following equation (for more details see supplementary information):

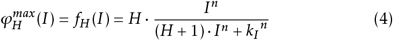

Here, *H* represents the maximum possible expression for the circuit, *I* is the molecular inducer concentration, *k*_*I*_ is related to the affinity for *I*, and *n* is the Hill coefficient that expresses the cooperativity of the response to *I*.

Now we will use the values of *f*_*i*_ (*t*)obtained from the maximum values of the heterologous fraction, 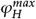, as described above, to test if they follow a dose-response profile in the shape of a typical Hill function. First, we verified by SDS PAGE analysis that even for the highest GFP overexpression, *φ*_*H*_ represented less than 15% of total biomass (Figure 5S); therefore, we infer that in most cases the coefficient (*H* + 1) can be approximated to 1, thus resulting in a typical Hill function.

Importantly, because in this kind of titration experiment, the curves represent the behaviour of the whole cell, *k*_*I*_ and *n* will not strictly correspond to the dissociation constant and the molecular cooperativity of the binding reaction between the transcription factor and the inducer.

For the DE3 expression system, the expression fractions, derived from experimental 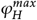values, related to the inducer concentration according to a Hill function. We achieved excellent fits in five different cell contexts, including three distinct *E. coli* strains with and without an additional LacI repressor (Figure 3). The parameters obtained from these fits are listed in Table 1. In contrast, extracting normalised fluorescence values at the time of maximum specific growth rate did not yield a Hill function in any of the five cases analysed. This is likely because the inducer (IPTG) is not yet equilibrated with the regulatory machinery at this point (which will be shown by the non-linear regions in the plots of Figure 4). Therefore, observations made at the time of maximum specific growth rate may not always accurately represent the behaviour of the biosensor regulatory circuit.

**Table 1.** Hill function parameters describing the biosensors responses. The Hill function was fitted to the values of the expression fraction plotted as a function of inducer concentration (IPTG, cumate, or HgBr_2_). The expression fractions were determined through two methods: Maxima, each fraction of expression is measured as equal to the heterologous protein fraction at its maximum value,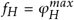. *Slopes*, each fraction of expression is determined by measuring the slope within the linear interval of the specific production rate vs specific growth rate plot,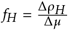.

| Sensor | Strain | $H(flu/OD_{600})$ | | $k_I (\mu M)$ | | $n$ | |
| --- | --- | --- | --- | --- | --- | --- | --- |
|  |  | Maxima | Slopes | Maxima | Slopes | Maxima | Slopes |
| IPTG | BL21(DE3) + LacI<sup>OV</sup> | 9784 ± 191 | 8457 ± 225 | 31.0 ± 3.7 | 35.6 ± 5.5 | 2.5 ± 0.3 | 2.5 ± 0.5 |
|  | BLR | 6148 ± 496 | 6668 ± 651 | 55.8 ± 12.9 | 58.0 ± 14.9 | 2.2 ± 0.3 | 2.7 ± 0.5 |
|  | BLR + LacI<sup>OV</sup> | 11473 ± 319 | 11080 ± 563 | 51.7 ± 5.1 | 67.7 ± 6.9 | 2.8 ± 0.3 | 4.4 ± 1.8 |
|  | MG1655(DE3) | 2072 ± 44 | 2846 ± 438 | 36.6 ± 4.1 | 56.3 ± 22.8 | 2.1 ± 0.2 | 2.0 ± 0.7 |
|  | MG1655(DE3) + LacI<sup>OV</sup> | 1116 ± 116 | 1747 ± 97 | 99.8 ± 4.5 | 93.6 ± 2.6 | 5.9 ± 1.4 | 6.6 ± 0.5 |
| Cumate | BL21(DE3) | 1170 ± 84 | 2965 ± 110 | 89.6 ± 13.4 | 79.7 ± 6.1 | 2.5 ± 0.2 | 4.3 ± 0.6 |
|  | BLR(DE3) | 989 ± 45 | 2390 ± 33 | 50.8 ± 5.3 | 33.5 ± 1.8 | 2.3 ± 0.2 | 4.0 ± 0.2 |
|  | MG1655(DE3) | 1539 ± 69 | 2575 ± 127 | 243 ± 17 | 208 ± 15.3 | 1.8 ± 0.2 | 3.2 ± 0.5 |
| Hg<sup>2+</sup> | NEB<sup>®</sup> Stable | - | 11722 ± 3130 | - | 554 ± 281 (nM) | - | 0.99 ± 0.1 |

**Figure 3.**
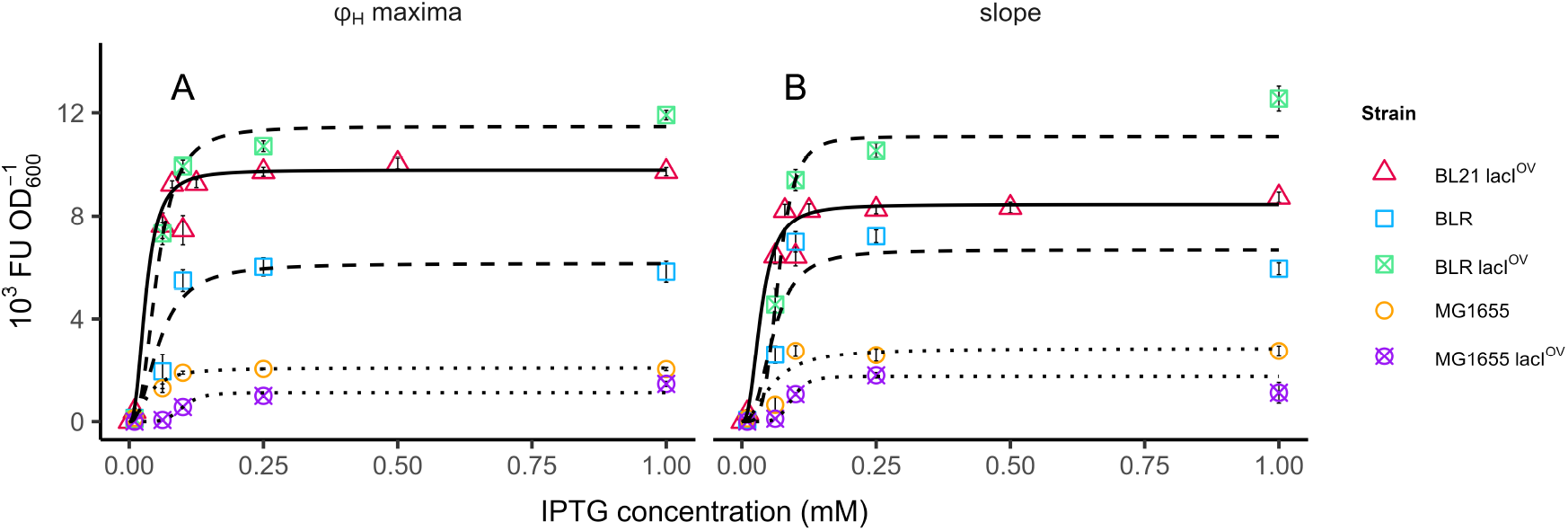
Expression fractions in response to the inducer, IPTG. A) Dose-reponse diagram made by plotting the maximum values of specific fluorescence (fluorescence over cellular density) obtained with the indicated IPTG concentration concentration in 5 different cellular contexts. B) Dose-reponse diagram made by plotting the slopes obtained in the linear regressions of the specific production vs specific growth graphs against its corresponding IPTG concentration. For both graphs the lines show the Hill function fitted for each data set. All strains bear the DE3 insert. Cells co-transformed with pACYCDuet for the additional expression of LacI are indicated as lacI^*OV*^. (N = 8 for BL21(DE3), and N = 4 for all other cases) ALT TEXT: Hill plots using slope method and maximum method for various strains with the DE3 system

**Figure 4.**
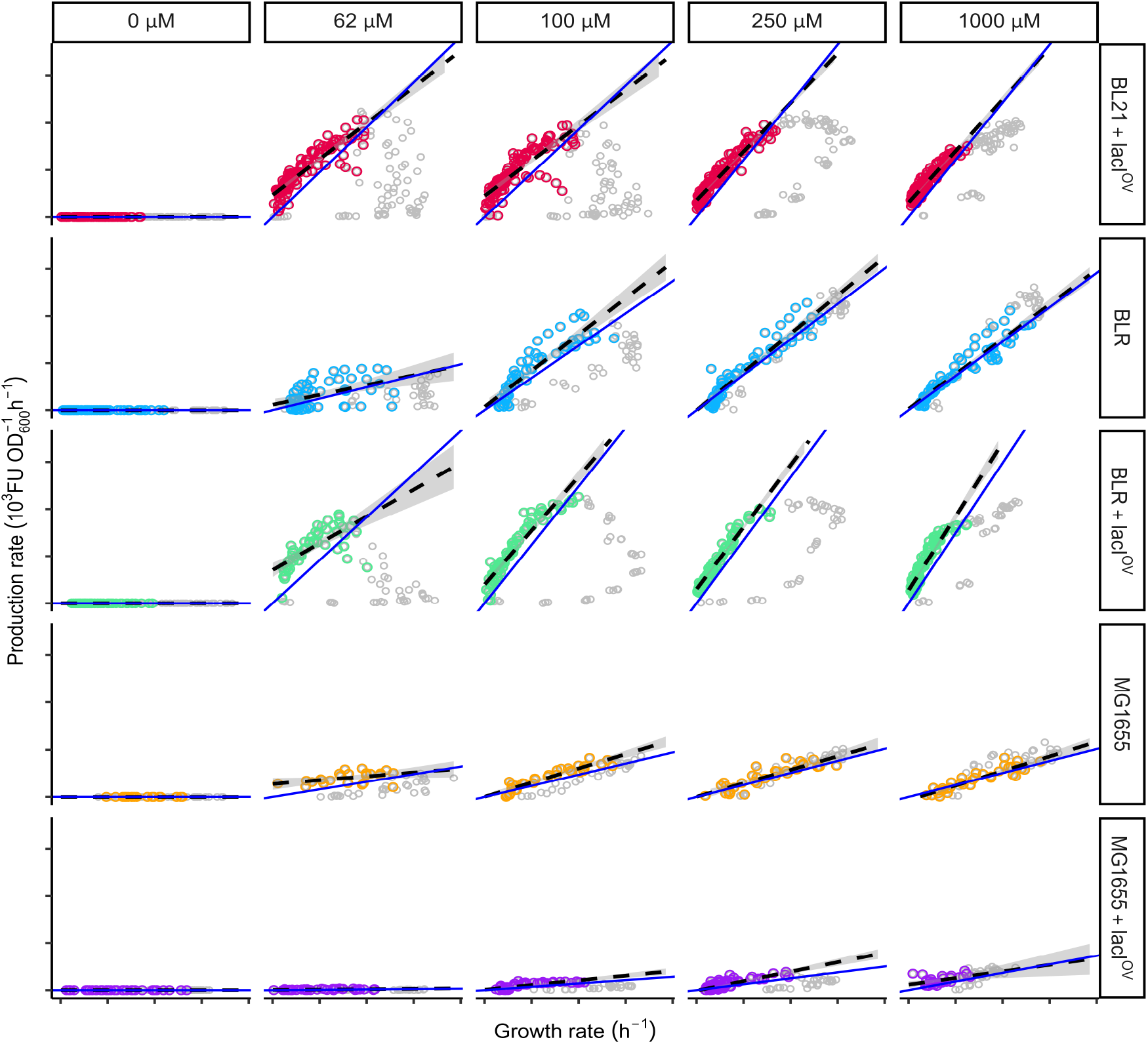
Expression fractions at different IPTG concentrations as obtained from two analysis methods. Experimental data taken after induction are directly plotted and then a time window is selected (colored dots) by excluding the earlier points (grey points) to be fitted to a linear regression (dashed line), maximizing R correlation coefficient as described in the main text. Each solid line represents a line function where the slope is set to be equal to the value of 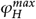 found at the indicated condition and strain. ALT TEXT: DE3 system Scatter plots of specific production rate vs specific growth rate. In each plot, a selected interval was used to make a linear regression. The blue line represents the maximum specific fluorescence represented as a slope.

### Finding the expression fraction during growth: The slope method

In the cases analysed, we observed that the maximum point for the heterologous fraction is reached close to or during the stationary phase. Because there are physiological changes in the onset of this phase, there is no guarantee that the biosynthesis fraction dedicated to our gene of interest will remain constant in other stages of the culture. Moreover, as stated previously, it is important for biosensor developers, especially in the field of metabolic engineering, to have a solid method to characterise heterologous expression during growth.

We aim to develop an effective analysis strategy by linking the expression fraction to the specific growth rate. By definition, the biosynthesis rate of the heterologous protein fraction, *ρ*_*H*_ (*t*), is the product of its expression fraction, *f*_*H*_ and the total protein biosynthesis rate, *ρ*(*t*):

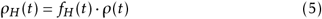

During exponential growth, the protein-to-total cellular-mass ratio remains relatively constant [50]. Therefore, we can confidently assume that the total protein biosynthesis rate remains linear to the specific growth rate, which implies that we can substitute *ρ*(*t*) with *µ*(*t*) without altering the proportionality of change. Since measuring the specific growth rate directly is simpler than determining the total biosynthesis rate of the cell, this approximation yields a convenient equation:

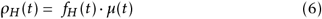

It follows that the *f*_*H*_ can be experimentally determined as equal to the slope in a plot of the specific production rate of the heterologous protein vs specific growth rate of the culture. As an approximation, we graphed the fluorescence specific production rate (*ρ*_*H*_) against the specific growth rate (*µ*) at all time points in Figure 4. Following induction, our initial data points from each culture show a curved line (grey dots) resembling a hook. This curved line transitions into a straighter segment (colored dots) that can be approximated by a linear relationship. We interpret this initial curvature as the time it takes for the inducer and the regulatory machinery to reach a steady state. The following straighter segment, on the other hand, represents a period with a constant slope or expression fraction (*f*_*H*_) of the heterologous protein. We used the resulting *f*_*H*_ values to generate a dose-response diagram and fit a Hill function, similar to the analysis performed with the values obtained from the 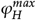 method (Figure 3).

Another important step is to investigate whether the *f*_*H*_ values obtained from 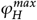 are similar to those obtained from the slopes. This would indicate that *f*_*H*_ remains constant throughout the interval in which the linear trend is maintained, including a significant portion of exponential growth and the beginning of the stationary phase. In Figure 4 (and Figure 1S) we present a visual comparison of the values *f*_*H*_ determined from the slope of the specific production versus the specific growth rate (dashed lines) and from the maximum heterologous fraction, 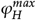 (solid lines). The similarity of the results obtained through these two different methods in the case of the DE3 system suggests that the expression fractions are indeed constant within these intervals, including the beginning of the stationary phase, where the values of 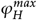 were identified.

Furthermore, it will be important to compare the Hill parameters (*H, k*_*I*_, *n*) obtained by fitting the *f*_*H*_ values from both the maximum method and the slope method. Table 1 presents the Hill parameters obtained by fitting these values *f*_*H*_ as a function of inducer concentration in different expression systems.

### Application on other biosensors (non T7RNAP)

The DE3 expression system depends on transcription driven by T7RNAP, which is suitable for various applications but somewhat restricted in terms of regulatory possibilities. In contrast, numerous transcription factors and other regulatory elements have been identified in relation to bacterial RNAP. To evaluate the broader applicability of our methods, we examined inducer titration experiments from two additional systems using transcription by *E. coli* RNAP.

First, a mercury biosensor was constructed based on the MerR transcription factor (complete characterisation in [37]). Upon ionic mercury titration, we observed that the reporter protein accumulated continuously, even during the stationary phase; therefore, we were unable to identify 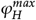. However, the specific production vs specific growth rate graphs (Figure 5) produced linear shapes. A linear region was detected in the plots and was approximated to a single slope that intersected at zero. A dose-response plot was plotted using the expression fractions obtained from these slopes and was fitted to a Hill function (Figure 4S). In addition, a cumate biosensor was built based on the constitutive expression of CymR repressor as seen in previous reports [38]. In the case of cultures induced with cumate, the reporter protein production decreased earlier than the arrest of growth, that is, before the onset of the stationary phase. Consequently, a 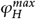 was readily identified. Also, we observed that the specific production vs specific growth rate graphs exhibited a very short lapse of nonlinear behaviour, indicating that the intracellular concentration of cumate equilibrates relatively rapidly. The linear lapse in the relation between specific production and specific growth rates showed an intersection different from zero on the specific growth rate axis (Figure 5 A), thus representing the period when specific production ceased while growth continued (*µ >* 0). Therefore, the regression in the cumate system must include the value of this intersect, thus implying that the equation for the specific production rate *ρ* is:

**Figure 5.**
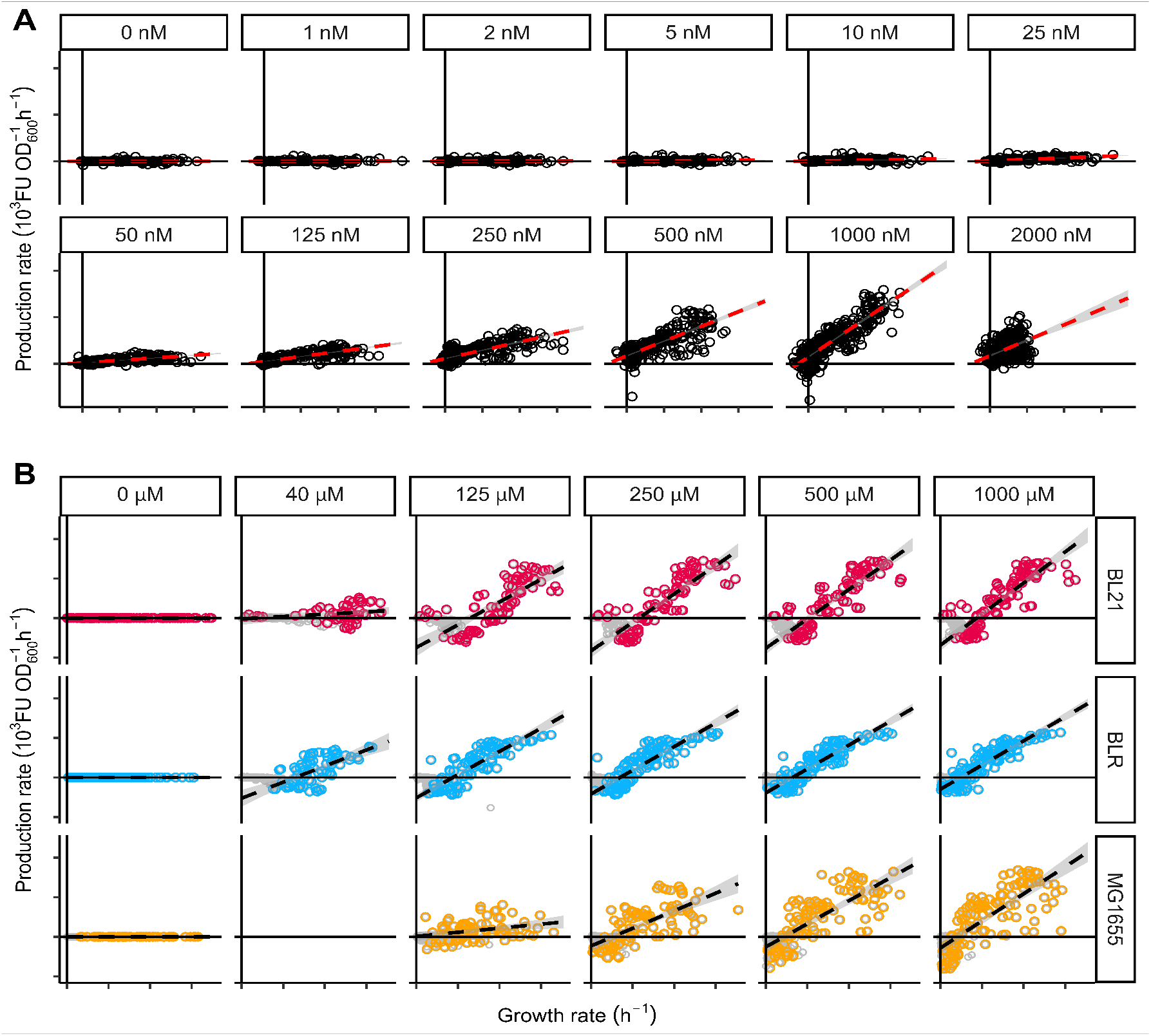
Expression fractions of biosensors at various inducer concentrations. A) Expression fractions of the MerR system in response to ionic mercury. Graphs show the RFP production upon induction with 0 to 2000 nM HgBr_2_, as indicated, in NEB^®^Stable cells. Dark dots are experimental data fitted to a linear regression (dashed line) whereas open circles correspond to early time points that are excluded to maximize the correlation coefficient as indicated in the main text. B) Expression fractions at different cumate concentrations in the CymR-cumate system. Graphs show the GFP production upon induction with 0 to 1000 *µ*M cumate, as indicated, in BL21(DE3), BLR(DE3) and MG1655(DE3) cultures transformed with the CymR-cumate system. Colored dots are experimental data fitted to a linear regression (dashed line) whereas grey dots correspond to early time points that are excluded to maximize *R*^2^ as indicated in the main text. The expression fractions here determined were used to graph dose-response diagrams (Figure 1S and 2S in Supplementary Material) and were fitted to Hill functions (parameters in Table 1). ALT TEXT: MerR system and CymR system Scatter plot of specific production rate vs specific growth rate. In each plot, a selected interval was used to make a linear regression.

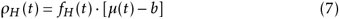

We anticipate that the intersection *f*_*H*_ (*t*) · *b* will be present in other biosensors when the maximum heterologous protein fraction 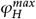 occurs at a positive specific growth rate (*µ >* 0). In systems like the CymR-cumate system, where the specific growth rate at the zero specific production rate intersection is significant (comparable to the slope value), it is no longer true that the maximum 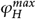 equals *f*_*H*_. Therefore, the regulatory parameters extracted from the dose-response plots will differ when using the 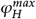 method compared to the slope method. For illustrative purposes, Figure 2S presents Hill plots generated by both methods. However, the expression fractions obtained from the slope method should be considered reliable.

## Discussion

This research tackles the challenge of standardising methods for characterising regulatory circuits and their modular components across various cellular contexts. The consistent behaviour of these parts is crucial for the long-term goal of designing predictable biological systems based on well-understood individual components. To this aim, we utilise conceptual tools from bacterial physiology to facilitate the characterisation of sensing circuits, particularly focussing on transcription factor-based biosensors.

The conceptualisation of the cell as a limited proteomic space introduced by Scott et al. [25] and expanded by subsequent kinetic models [32, 26] addresses the interdependence between the specific growth rate and the regulation of different sets of genes. Using the time derivative of the formal definition of a proteome fraction, the focus is shifted from protein yield to protein biosynthesis rate. Following the two alternative methods presented here, the experimental results readily produce the expression fraction, which represents a protein-specific production rate. By means of inducer titrations, the experimenters may obtain the expression fraction at various inducer concentrations, which are then summarised in a single set of Hill function parameters.

We first evaluated our approach by characterising the DE3 system in three distinct strains, considering both the presence and the absence of additional expression of the LacI repressor. Although the slope and maximum methods yield reasonably similar results, it is evident that these parameters are significantly influenced by the cellular context. For instance, the hooked shapes in the rate graphs in Figure 4 illustrate the time interval during which the expression fraction is not constant, presumably because the inducer has not yet reached equilibrium with the transcription factor. This lag period is shorter for higher concentrations of the inducer, as one would expect from a diffusion-dependent equilibration. Furthermore, it seems to be strain dependent and is associated with B strains exhibiting a higher apparent affinity for the inducer, as shown by a lower value of *k*_*I*_ than MG1655(DE3). These variations cannot be attributed to actual differences in the association reaction between the LacI protein and IPTG; therefore, these observations could be attributed to variations in IPTG permeability, considering the regulation of lactose permease (LacY) and the relative contribution of passive diffusion and active transport among these strains. A K-12 strain has been reported to depend on inducible LacY for the internalisation of IPTG [51, 52]. In agreement with this, when LacI overexpression was introduced in MG1655(DE3), *k*_*I*_ increased three or four times its value, which means that the apparent affinity of the system decreased, probably due to the reduced internalisation of IPTG caused by the decrease in LacY expression. This careful analysis of the rate plot and the Hill parameters in different contexts provides valuable observations specific to the cells where the system is tested. In future studies, we plan to evaluate and deepen into more biosensor cases, including the biosensors for mercury and cumate presented here, to see in what cases the Hill parameters are kept across various cells, and in what cases the variations are informative of strain-specific characteristics.

We evaluated the general applicability of the two analytical methods (maximum and slope) by analysing titration experiments from two additional biosensing systems. These systems detect cumate and mercury using CymR and MerR transcription factors, respectively. The maximal fractions method was not suitable for the MerR system because the reporter protein continuously accumulated, even after entering the stationary phase, preventing the detection of a clear maximum point 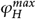. For the cumate biosensor, although the maximal fractions were readily identifiable, the presence of an intercept implies that 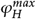 does not equal the expression fraction. This discrepancy explains the inconsistencies between the Hill parameters obtained for the CymR system from the maximum and slope methods. Despite these limitations with the maximum method, the slope method remained a reliable approach for both the CymR and MerR systems. The extended linear regions observed in the rates plots indicated the suitability of these systems for analysis using the slope method. Furthermore, the brief non-linear phase suggests a faster equilibration with their specific inducers compared to the slower equilibration observed with IPTG.

In addition to analysing their applicability conditions, we also consider the limitations of each method that should be acknowledged for future uses. The maximum fraction method relies on a single data point, often obtained during the stationary phase. This approach does not provide information on the interval of time at which the expression fraction thus determined is valid. Furthermore, cellular functions are down-regulated during the stationary phase, potentially compromising the suitability of the method for assessing the biotechnological applications of a system despite a high apparent heterologous expression level. In the case of the slope method, it employs a linear regression in a time window of constant slope in the data, thereby overcoming this limitation of the maximum method. However, the ranges of linear correlations must be identified for each analysed culture condition, which involves a potential source of error. In this method, the selection of points for the regression analysis might be subjective or insufficient if dealing with a small number of data points. To address this limitation, we propose defining strict data requirements and implementing an algorithm for a more objective determination of the linear range. Other improvements will arise from better protein measurement accuracy. Here, we approximated the reporter proteome fraction, *φ*_*H*_, simply normalising the fluorescence by OD_600_. Better measurements can be made by complementary techniques, such as relating the registered fluorescence intensity to specific amounts of reporter protein through a calibration curve and MS-supported proteomic approaches.

In broad terms, this work presents methods and concepts designed to aid researchers in applying principles from advanced metabolic studies to characterise new regulatory elements and develop novel biosensor circuits. For example, a significant challenge in biocircuitry design is the lack of formal evaluation of orthogonality between heterologous gene expression and native functions [3, 53]. Existing methods often lack standardisation and rely on comprehensive and expensive techniques such as RNAseq and ribosome profiling [54, 55, 56, 57]. Alternatively, researchers employ specific, hypotheses-driven tests to identify interference between regulatory components [58, 59, 60, 61]. In contrast, the framework presented here offers a relatively simple approach to assess how the regulations of native functions, composed of expression fractions *f*_*C*_ (nutrient fixators) and *f*_*R*_ (biomass producers), are influenced by *f*_*H*_ (heterologous expression). For example, researchers can manipulate the carbon sources in the culture to observe changes in *f*_*C*_ [25, 62, 32]. Alternatively, physical or biochemical stress mediated by ppGpp can be used to monitor changes in *f*_*R*_ [26, 63, 64]. These manipulations will activate distinct cellular regulatory mechanisms and, if a novel heterologous circuit affects these mechanisms, a dose-response relationship should emerge, with increased heterologous expression correlating with heightened interference. This approach could reveal interactions between native metabolic regulations or stress responses and the heterologous gene(s). In this way, the concepts of proteome fractions and expression fractions offer a more nuanced approach to evaluate reporter protein production compared to simple normalisation by culture density or growth rate. These methods remain cost-effective and convenient, potentially encouraging wider adoption of a systems-wide approach in the characterisation of biomolecular parts for synthetic biology and improving reproducibility across laboratories.

## Author information

### Author contribution

**Nicolás A. Vaccari** Conceptualised and developed the model, designed and cloned the IPTG biosensor system, performed titration experiments, performed data analysis and wrote the article. **Dahlin Zevallos-Aliaga** - Designed and cloned the biosensors for mercury and cumate and performed their titration assays. **Tom Peeters** - Provided critical feedback on the design and testing of the mercury biosensor. **Daniel G. Guerra** - Served as project supervisor, providing guidance and critical feedback for experimental design, modelling, and data analysis, and wrote the article.

### Notes

The authors declare that they have no competing financial interests.

## Material availability statement

The sequences of the genetic circuits designed for this study are available on GenBank: IPTG biosensor (GenBank: PQ015608), and the cumate biosensor (GenBank: PQ010742). The sequence of the mercury biosensor is described in [37]. All plasmids used in this study will be shared upon request to the corresponding author.

## Data availability statement

Datasets available on FigShare: https://doi.org/10.6084/m9.figshare.263147213.vM1

## Acknowledgements

This work was supported by CONCYTEC – PROCIENCIA - Contrato 170-2020. The IPTG and cumate biosensors were developed and characterised thanks to the support of Fondo Nacional de Desarrollo Científico, Tecnológico y de Innovación Tecnológica – FONDECYT / PROCIENCIA, through the program EF-041 - Proyectos de Investigación básica y aplicada. Grant number CONV-000114- 2015-FONDECYT-DE. The development of the mercury biosensor was funded by VLIR-UOS through the South Initiative grant code PE2020SIN292B122.

## Supplementary information

Supporting Experimental Information: Additional experimental results showing dose-response diagrams for the MerR and CymR systems, total protein analysis by SDS-PAGE for all strains containing the DE3 system expressing GFP, and total protein quantification for induced and uninduced bacteria. Supporting Mathematical Framework: Detailed definitions, step-by-step derivations, and algebraic arrangements.

