## Supplementary material for "Biosensors Characterization: Formal methods from the Perspective of Proteome Fractions": sup1

### Biosensors Characterization:

### SUPPLEMENTARY INFORMATION: SUPPORTING EXPERIMENTAL INFORMATION

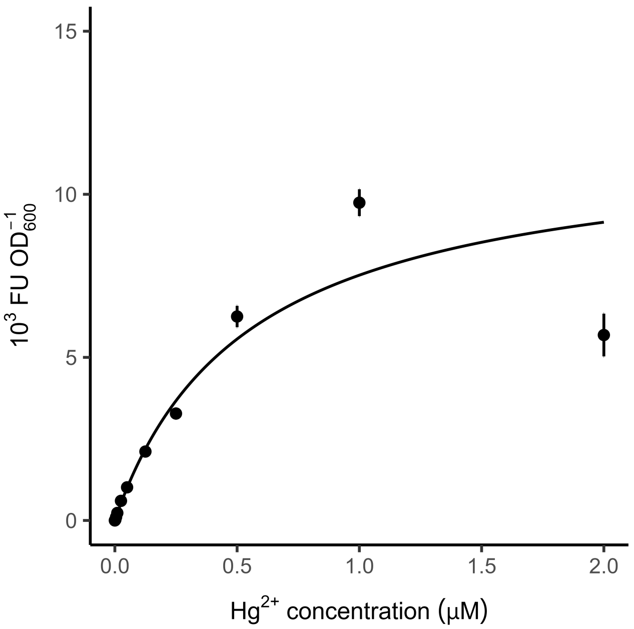


Figure 1: **Dose-response diagram for a mercury biosensor.** Dose-reponse diagram made by plotting the expression fractions obtained at various mercury concentrations. The expression fractions correspond to the slope values of the linear regressions shown in the specific production vs specific growth graphs (Figure 5A in the main text). The lines indicate the fitting to a Hill function. ALT TEXT: Hill plots using the expression fractions obtained from the slope method for the MerR system

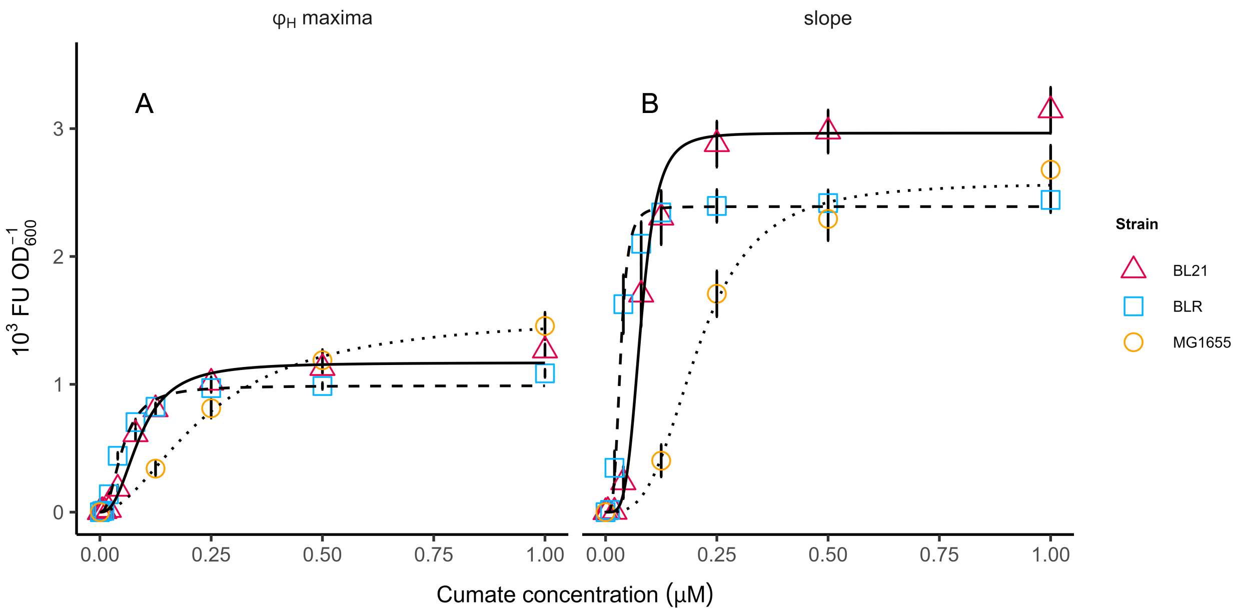


Figure 2: **Dose-response diagrams for a cumate biosensor** A) Dose-reponse diagram made by plotting the maximum values of specific fluorescence ($\varphi_H^{max}$) obtained with the indicated cumate concentration in three different cellular contexts, BL21(DE3), BLR(DE3) and MG1655(DE3). B) Dose-reponse diagram made by plotting the slope values of the linear regressions of the specific production vs specific growth graphs obtained with the indicated cumate concentration (Figure 5B in the main text). For both graphs the lines show the Hill function fitted for each data set. ALT TEXT: Hill plots using the expression fractions obtained from the maximum method and the slope method for the CymR system

.

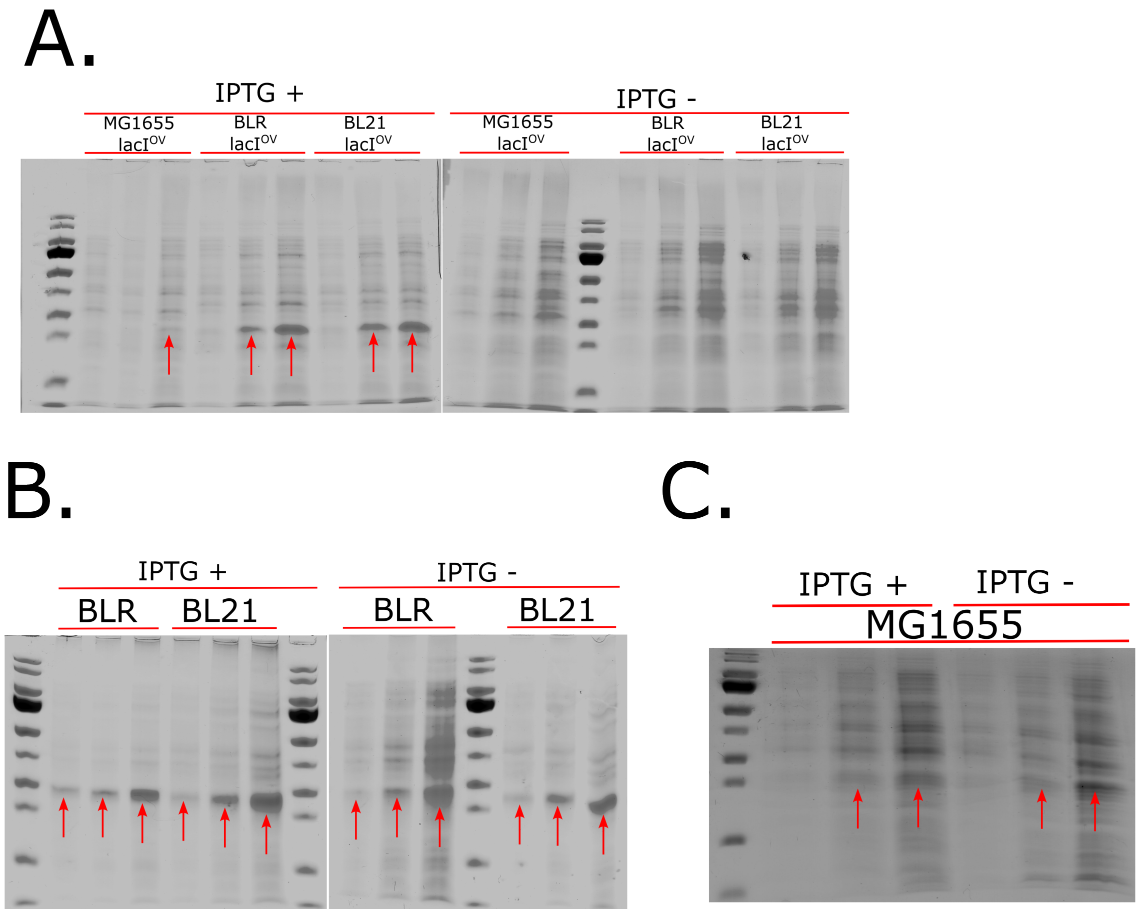


Figure 3: **Total protein from cells transformed with pUC-T7-GFP or pUC-T7-GFP and pACYCDuet with and without IPTG induction.** Total proteins from cultures of similar densities were solubilised with 8M urea and analysed through SDS-PAGE as described in Methods section. Red arrows indicate the bands corresponding to GFP. Image analysis determined that B strains could accumulate GFP up to roughly 15% of the total protein mass. In the case of MG1655(DE3), the cell accumulated a GFP amount equivalent to approximately 1% of its total protein mass. ALT TEXT: Total protein from induced and uninduced cells analysed through electroporesis to asses the fraction of the total proteome represented by GFP
