## Supplementary material for "Biosensors Characterization: Formal methods from the Perspective of Proteome Fractions": sup2

### Biosensors Characterization:

### SUPPLEMENTARY INFORMATION: SUPPORTING MATHEMATICAL FRAMEWORK

#### Initial definitions

##### Proteome fractions

A particular set of proteins $i$, will be defined here as a proteome fraction, $\varphi_i$, by dividing the total mass of said group of proteins, $m_i$, over the mass of the complete proteome, $m$:

$$\varphi_i(t) = \frac{m_i(t)}{m(t)} \tag{1}$$

where:

$$m(t) = \sum_i m_i(t) \tag{2}$$

**Proteome fluxes & expression fraction**

To study how $\varphi_i$ evolves over time, we need to define the anabolic flux, which is the velocity at which the cell produces proteins. We define this as the product of the total anabolic rate $\rho$ and the mass of the total proteome $m$:

$$\frac{dm(t)}{dt} = \rho(t) \cdot m(t) \tag{3}$$

Also, as mentioned in the main text, the total anabolic flux can be divided into sub-fluxes that produce the different protein groups:

$$\frac{dm_i(t)}{dt} = f_i(t) \cdot \frac{dm(t)}{dt} \tag{4}$$

$$\frac{dm_i(t)}{dt} = f_i(t) \cdot \rho(t) \cdot m(t) \tag{5}$$

Where $f_i$ is the expression fraction, which represents the fraction of the total anabolic flux dedicated to producing a set $i$ of proteins. From equation (2), (3) and (5) we can deduce that:

$$\rho(t) = \sum_i f_i(t) \cdot \rho(t) \tag{6}$$

Here, we will define the specific production rate of protein $i$, or $\rho_i$, as:

$$\rho_i(t) = \frac{1}{m(t)} \cdot \frac{dm_i(t)}{dt} \tag{7}$$

Then, we can define each corresponding production rate for each of the protein sets using equations 5 and 7 to obtain the production rate:

$$\rho_i(t) = f_i(t) \cdot \rho(t) \tag{8}$$

#### Biosynthesis allocation theorem

To study proteome fractions and their change over time, we start by unfolding the derivative of $\varphi_i$ as the derivation of a division:

$$\frac{d\varphi_i(t)}{dt} = \frac{d\left(\frac{m_i(t)}{m(t)}\right)}{dt} \tag{9}$$

Expanding and resolving:

$$= \frac{m(t) \cdot \frac{dm_i(t)}{dt} - m_i(t) \cdot \frac{dm(t)}{dt}}{m^2(t)}$$

$$= \frac{m(t) \cdot [f_i(t) \cdot \rho(t)] \cdot m(t) - m_i(t) \cdot \rho(t) \cdot m(t)}{m^2(t)}$$

$$= \rho(t) \cdot \left(f_i(t) - \frac{m_i(t)}{m(t)}\right)$$

$$\frac{d\varphi_i(t)}{dt} = \rho(t) \cdot [f_i(t) - \varphi_i(t)] \tag{10}$$

Hence, the evolution of $\varphi_i$ depends on the anabolic rate $\rho$ and the difference between $f_i$ and $\varphi_i$. This expression is very similar in form to that of potential-induced currents, where $\frac{1}{\rho}$ can be interpreted as resistance, and the difference between $f_i$ and $\varphi_i$ can be understood as a sort of differential potential, where $f_i$ represents the fraction value of the proteome intended by the cell and $\varphi_i$ represents the value that it actually has. In general, this analysis provides a framework for an easy understanding of the dynamics of proteome fractions.

#### Total anabolic rate and specific growth rate

Because measuring the anabolic rate can be challenging, we will look for a particular and more useful form of the theorem to express it as a function of the growth rate. When the ratio of protein mass to biomass is constant, the following proportionality is true:

$$\frac{dm(t)}{dt} \propto \frac{db(t)}{dt} \tag{11}$$

where $b$ is biomass and follows the accumulation equation:

$$\frac{db(t)}{dt} = \mu(t) \cdot b(t) \tag{12}$$

Here $\mu$ is the growth rate, a classical parameter that can be easily measured as the change in optical density over time. Using Equations 3, 9, and 10 we obtain that as long as the protein to biomass ratio remains constant:

$$\mu(t) = \rho(t) \tag{13}$$

From experimental measures, it is known that this condition is met during the exponential phase of bacterial growth[?] . Rearranging the equation (10), for this particular case we obtain:

$$\frac{d\varphi_i(t)}{dt} = \mu(t) \cdot [f_i(t) - \varphi_i(t)] \tag{14}$$

#### Regulation function and expression fraction

##### Definitions

In general, it is possible to represent the rate of expression of a set $i$ of proteins as:

$$\widehat{\rho_i}(t) = r_i(t) \cdot \rho(t) \tag{15}$$

In this equation, $r_i$ can represent a repression term, induction term, or constitutive term. We will refer to $r_i$ as the regulation function because it modifies the behaviour of the protein production flux for a specific set of proteins. Here, we will provide functions to represent three different modes of regulation.

The first case considers the production of a set of proteins $i$ that is repressed by a known concentration of a transcription factor $TF$, with a constant of repression $k_{TF}$. We can model this regulated production flux as follows:

$$\widehat{\rho_1}(t) = \theta_1 \cdot \frac{k_{TF}}{TF(t) + k_{TF}} \cdot \rho \tag{16}$$

The second case represents the expression as induced by a known concentration of an inducer $I$, with an induction constant $k_I$:

$$\widehat{\rho_2}(t) = \theta_2 \cdot \frac{I(t)}{I(t) + k_I} \cdot \rho(t) \tag{17}$$

In both equations, $\theta$ is a coefficient that represents the maximum activation capacity of the regulatory circuit that governs the set of proteins. In other words, it indicates what fraction of total protein production is directed toward this set of proteins when the system is perfectly derepressed ($TF = 0$) or perfectly induced ($I \to \infty$).

Finally, a third case is a constitutive mode of expression, where there is no repressor nor inducer modifying the production flux. This mode of regulation is represented only by the maximum activation capacity coefficient $\theta$:

$$\widehat{\rho_3}(t) = \theta_3 \cdot \rho(t) \tag{18}$$

In all cases, the regulation function, $r_i$, is an abstraction that only represents the behaviour of the overall regulatory mechanism of a set of proteins. This regulatory function can be very complex or relatively simple, as in the three cases defined above. Compiling various proteins into a single set and generalising a unified $r_i$ for them may require the inclusion of several environmental factors and intracellular transduction pathways.

By definition, the sum of all fluxes of all proteome fractions must be equal to the total anabolic flux. This means that the sum of all regulation functions can be normalised.

$$\frac{r_1(t)\cdot\rho(t)+r_2(t)\cdot\rho(t)+r_3(t)\cdot\rho(t)+\ldots+r_n(t)\cdot\rho(t)}{[r_1(t)+r_2(t)+r_3(t)+\cdots+r_n(t)]}=\rho(t) \tag{19}$$

From this expression, we deduce that the production rate of a proteome fraction, $\varphi_i$, is:

$$\rho_i(t)=\frac{r_i(t)}{\sum_j r_j(t)}\cdot\rho(t) \tag{20}$$

Recurring to equation (5) and using this last equation, we finally define the expression fraction $f_i$, which, as mentioned before, represents the fraction of the total anabolic flux dedicated to produce a set $i$ of proteins, as:

$$f_i(t)=\frac{r_i(t)}{\sum_j r_j(t)} \tag{21}$$

**Introducing a heterologous protein**

The T7 expression system is a convenient regulatory circuit for studying heterologous genetic circuits. It provides a well-known regulation mechanism for the protein being expressed under its control. In addition, it is a system that includes its own RNA polymerase, which is not directly influenced by any metabolite or additional transcription factor. Finally, when activated by IPTG, its regulatory function can be considered constant because IPTG is a nonmetabolisable ligand whose intracellular concentration, once it reaches equilibrium, remains stable:

$$r_H(t)=H\cdot\frac{I}{I+k_i} \tag{22}$$

Here, $I$ is the concentration of an inducer, $H$ is the maximum expression activation number for the heterologous circuit, and $k_i$ is its induction constant. Following the theorem expressed in Equation (10), we obtain the change in time of the proteome fraction represented by the heterologous protein during the exponential phase:

$$\frac{d\varphi_H(t)}{dt} = \rho(t) \cdot \left( \frac{H \cdot \frac{I}{I+k_i}}{1 + H \cdot \frac{I}{I+k_i}} - \varphi_H(t) \right) \tag{23}$$

Future models could include a function for the internalisation of a nonconstant inducer $I(t)$, or its production and consumption within the cell.
